# Unmasking Glycoforms: Lectin-Based Profiling and Functional Implications of Targeted Glycosylation Knockouts in CHO Cells

**DOI:** 10.64898/2026.05.13.724788

**Authors:** Cristina Abascal Ruiz, Sheryl Li Yan Lim, Jacobus Brink, Sara Carillo, Eoin Casey, Jonathan Bones, Ioscani Jiménez del Val

**Author notes:** These authors contributed equally to the work (Cristina Abascal Ruiz, Sheryl Li Yan Lim).

## Abstract

Monoclonal antibody (mAb) glycosylation is a critical quality attribute that is difficult to rationally engineer and rapidly assess during cell line development. Here, we investigate whether cell-surface glycosylation can serve as a predictive indicator of mAb product glycosylation following targeted glycogene engineering in CHO cells. Five key glycogenes (COSMC, FUT8, B4GALT1, ST3GAL4, ST6GAL1) were investigated in two mAb-producing CHO cell lines. Product glycan analysis revealed consistent, gene-specific effects across hosts, including loss of core fucosylation, and tuneable galactosylation and sialylation. Lectin-based surface profiling reliably reflected product outcomes for COSMC and FUT8 modifications but showed limited predictive power for galactosylation and α2,3-sialylation, highlighting glycosylation pathway redundancy and context dependence. This study provides the first systematic, cross-cell line evaluation of lectin-based cell-surface glycan profiling as a predictor of mAb product glycosylation, establishing its practical utility and inherent limitations for CHO glycoengineering workflows.

**Graphical abstract:** 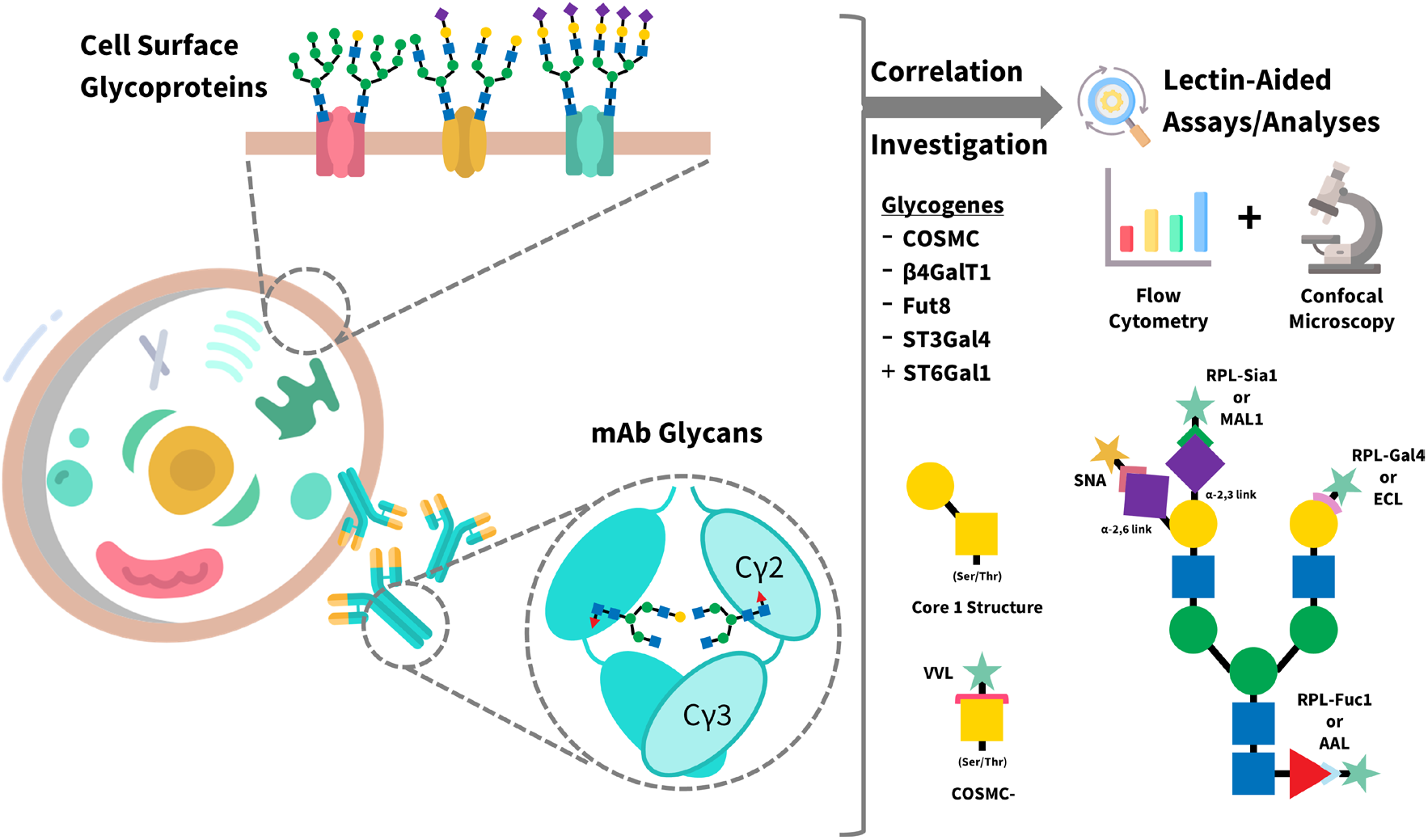

## Introduction

Monoclonal antibodies (mAbs) constitute the largest and most rapidly expanding class of biopharmaceuticals, with clinical efficacy, pharmacokinetics, and safety profiles that are tightly linked to glycosylation of the Fc and Fab domains [1–3]. In particular, variations in core fucosylation, galactosylation, and sialylation can profoundly influence antibody-dependent cellular cytotoxicity (ADCC), complement activation, serum half-life, and immunogenicity [4–8]. As a result, glycosylation is recognised as a critical quality attribute (CQA) of therapeutic antibodies, and substantial effort has been devoted to engineering host cell lines, most commonly Chinese hamster ovary (CHO) cells, to achieve defined and reproducible glycan profiles [9].

Despite major advances in genome editing and pathway engineering, the development of glycoengineered CHO cell lines remains time- and resource-intensive [10]. A persistent challenge lies in the rapid identification and characterisation of engineered clones with the desired product glycosylation phenotype. Direct analysis of secreted antibody glycans typically requires cell expansion, protein purification, and specialised analytical workflows, creating bottlenecks during early-stage cell line development. Consequently, there is considerable interest in proxy readouts that could enable faster screening and triage of engineered populations. One such approach is the assessment of cell-surface glycosylation, which reflects the activity of glycosylation pathways within the secretory system and can be interrogated rapidly using lectins [11].

Lectin-based profiling has been widely applied to monitor global changes in cell-surface glycosylation and, in selected cases, to enrich engineered populations with flow cytometry [12]. However, the extent to which cell-surface glycan phenotypes reliably reflect secreted mAb glycosylation remains unclear and appears to be highly context dependent. Glycosylation of membrane proteins and secreted antibodies occurs within shared Golgi compartments, yet their glycomes are shaped by distinct substrate specificities, trafficking routes, and enzyme redundancies [13]. Establishing when surface glycosylation can serve as a meaningful surrogate for product glycosylation, and when it cannot, remains an open question for CHO cell engineering.

In this study, we systematically examine this relationship by focusing on five glycogenes that are central to antibody glycosylation and are, in principle, amenable to lectin-based interrogation at the cell surface: COSMC, FUT8, β4GALT1, ST3GAL4, and ST6GAL1. These genes span key steps in O-glycan extension, core fucosylation, galactosylation, and sialylation, and include both loss-of-function and gain-of-function perturbations. To assess the robustness of observed phenotypes, these modifications (knockouts and/or knock-ins) were introduced into two distinct mAb-producing CHO cell lines, CHO VRC01 and NISTCHO, enabling evaluation of consistency across closely related but non-identical CHO backgrounds.

By integrating product glycan analysis with lectin-based flow cytometry and confocal microscopy, this work aims to define the strengths and limitations of cell-surface glycosylation profiling as a tool for glycoengineered cell line characterisation. Beyond its utility as a screening strategy, we assess the functional implications of surface glycan phenotypes and their interpretability in the context of enzyme redundancy, pathway architecture, and host-cell background. Together, these findings provide a framework for the rational use of lectin-based assays in CHO glycoengineering workflows and clarify their role in accelerating early-stage cell line development.

## Results

### Generation of Glycogene-Engineered CHO Cell Lines

Targeted perturbations of five glycogenes (COSMC, FUT8, β4GALT1, ST3GAL4, and ST6GAL1) were introduced into two mAb-producing CHO cell lines, CHO VRC01 and NISTCHO. These modifications included gene knockouts (COSMC, FUT8, β4GALT1, ST3GAL4) and overexpression strategies (ST6GAL1), as listed in Figure 1a. Clonal cell lines were established and verified by non-deletion/deletion PCR (Supplementary Figure S1 and Table S2), after which antibody products were harvested for glycan analysis.

**Figure 1:**
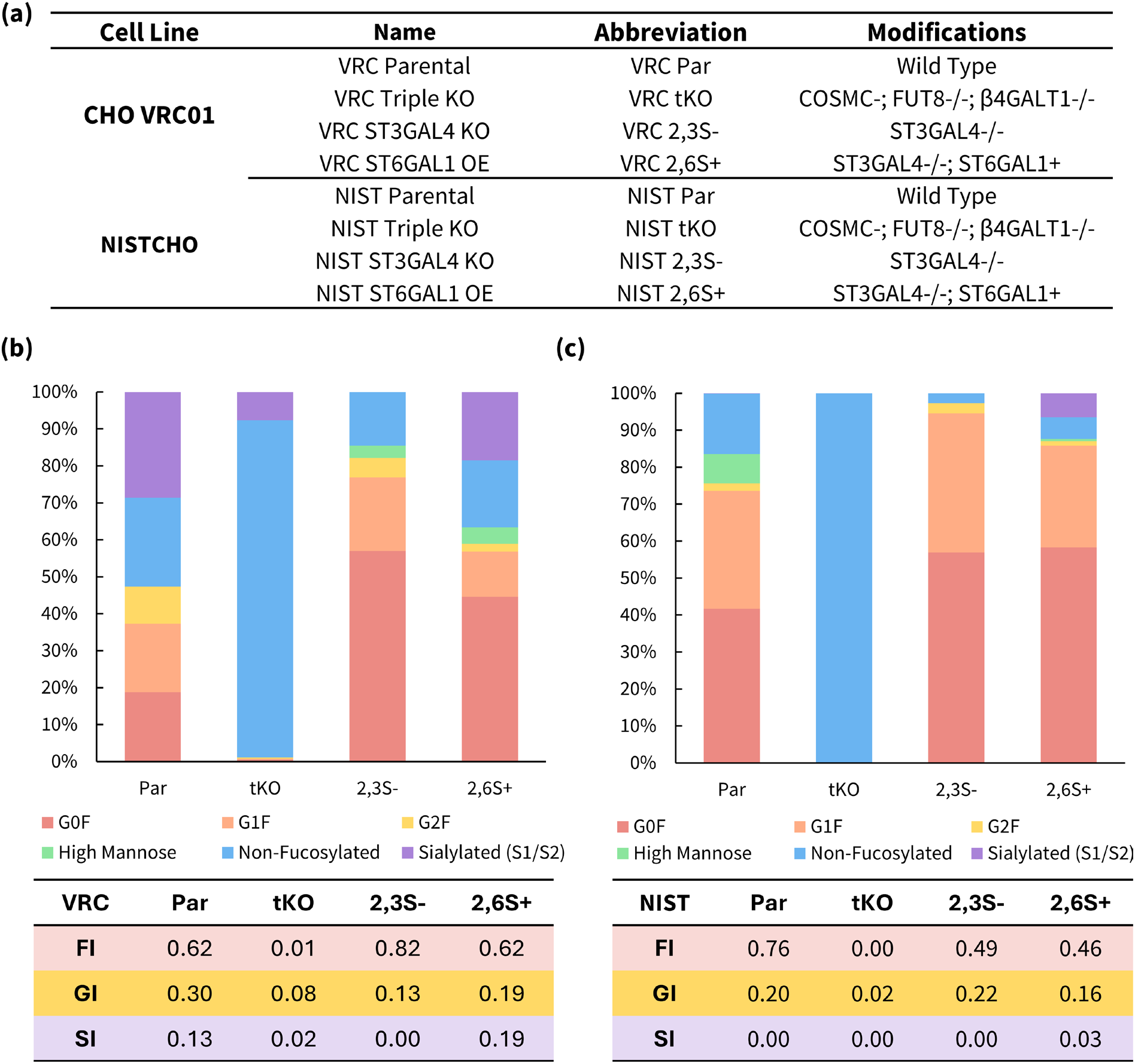
Product glycan distributions and glycosylation indices of engineered CHO cells. (a) Nomenclature of cell lines in the study; (b) CHO VRC01 cell lines; and (c) NISTCHO cell lines.

### Product Glycosylation Outcomes Following Targeted Glycogene Engineering

Product glycan analysis revealed distinct and gene-specific effects that were consistent across both CHO backgrounds. FUT8 knockout led to complete abrogation of core fucosylation on mAb glycans, as expected, while preserving overall glycan complexity.

Manipulation of galactosylation and sialylation pathways produced more nuanced outcomes. β4GALT1 knockout resulted in near-complete loss of mAb galactosylation and ST3GAL4 knockout eliminated detectable α2,3-sialylated species without affecting overall glycan occupancy, while ST6GAL1 overexpression introduced substantial α2,6-sialylation, a modification largely absent in wild-type CHO-derived antibodies. It is also important to note that there is limited native sialylation in the NISTmAb, which explains the modest increase (~6–7% of the total population) in the ST6GAL1 overexpression efforts in the NISTCHO cells. Together, these data establish clear and reproducible product glycosylation phenotypes for each targeted glycogene in both host backgrounds (Figures 1b and c).

### Cell-Surface Glycosylation Profiling by Lectin-Based Analysis

Having established product-level outcomes, we next assessed whether these engineered modifications were reflected at the cell surface using lectin-based profiling. Initial optimisation identified day 3 of culture as the most stable time point for surface glycan analysis, avoiding confounding effects associated with late-stage culture stress. Although this precedes the final antibody harvest (day 7/8), early time-point analysis was intentionally selected to enable rapid screening and cell enrichment via FACS during early-stage cell line development. At this stage, cell-surface glycosylation is more stable and less influenced by culture-induced artefacts, providing a reliable readout of the underlying glycoengineering modifications.

Lectin-assisted flow cytometry revealed strong concordance between surface glycan abundance and product phenotypes for glycogenes exhibiting binary functional behaviour. In particular, FUT8 knockout cells showed a robust reduction in fucose-specific lectin binding, confirming loss of core fucosylation at the cell surface, while COSMC knockout cells exhibited markedly increased *Vicia villosa* lectin (VVL) binding, consistent with GalNAc exposure on the cell surface [14]. These effects were consistent across both CHO VRC01 and NISTCHO backgrounds (Figures 2a, b, g).

**Figure 2:**
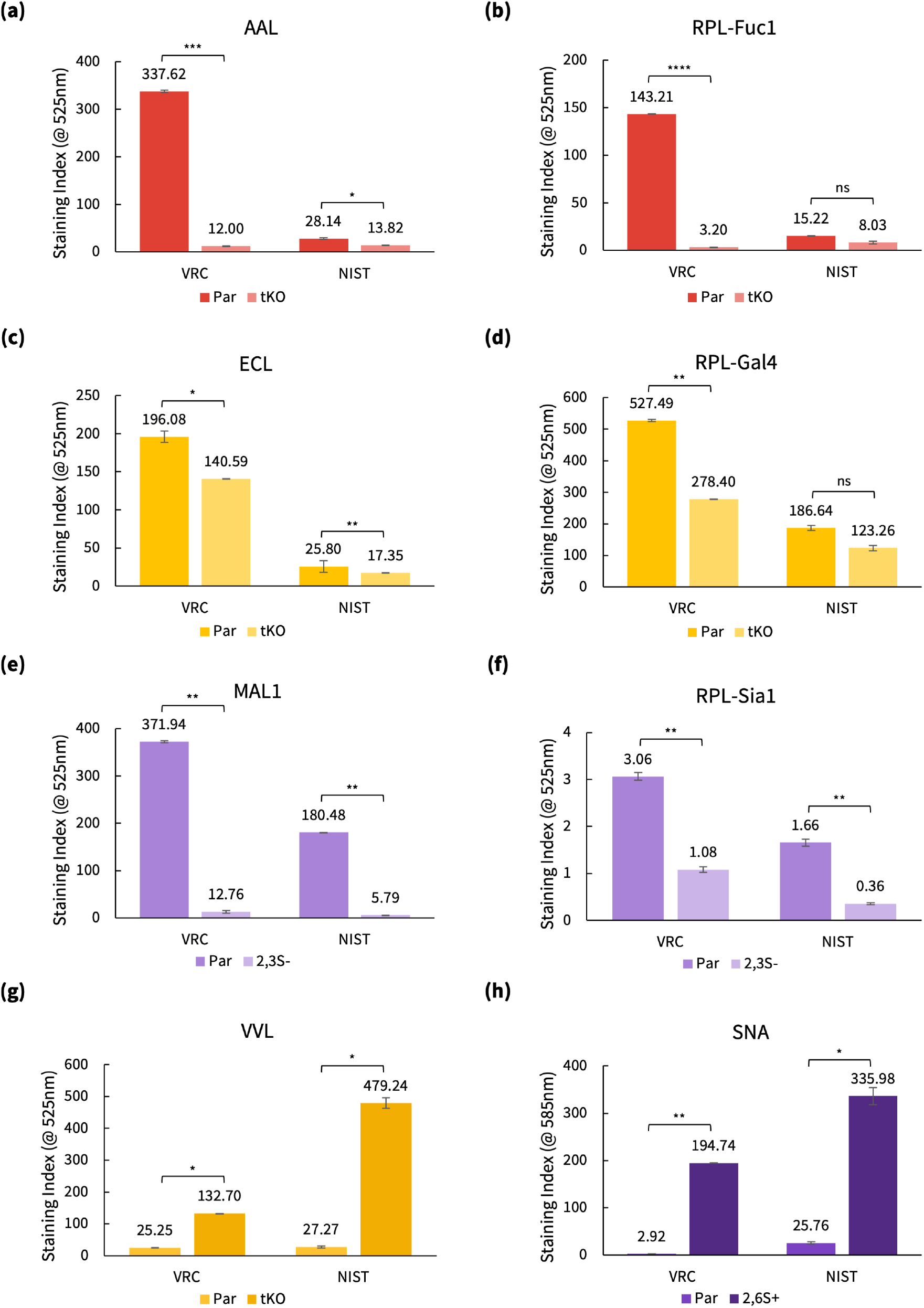
Staining indices of lectin-aided flow cytometry of engineered CHO cells. (a) AAL staining for FUT8−/−in tKO cells; (b) RPL-Fuc1 staining for FUT8−/−in tKO cells; (c) ECL staining for β4GALT1−/−in tKO cells; (d) RPL-Gal4 staining for β4GALT1−/−in tKO cells; (e) MAL1 staining for ST3GAL4−/−; (f) RPL-Sia1 staining for ST3GAL4−/−; (g) VVL staining for COSMC-; and (h) SNA staining for ST6GAL1 overexpression.

Confocal microscopy of lectin-stained cells supported these findings qualitatively. However, for select conditions, the magnitude of distinction between parental and engineered cell lines was less pronounced than that observed by flow cytometry. This was most evident for *Maackia amurensis* lectin I (MAL1) staining of parental versus ST3GAL4 knockout cells and for VVL staining of COSMC knockout cells in the NISTCHO background (Figures 3a and b). While lectin binding remained detectable at the cell surface by imaging, population-averaged flow cytometry measurements revealed a clear reduction or enhancement in total surface lectin binding, respectively.

**Figure 3:**
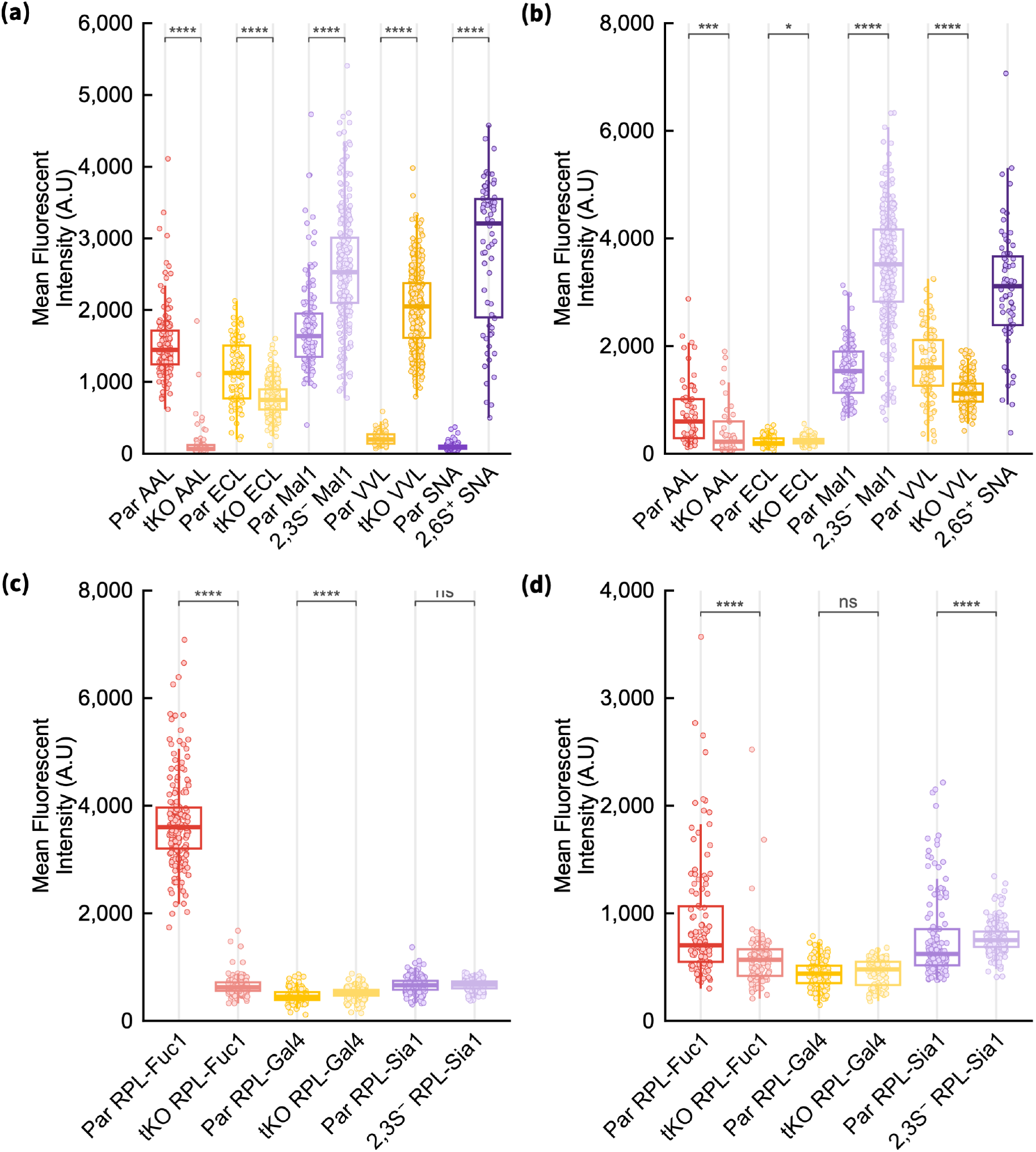
Mean fluorescence intensities (MFI) of lectin-stained cell samples from confocal microscopy analysis to determine and correlate cell surface with mAb glycan profiles. (a)CHO VRC01 cell lines stained with plant-derived lectins; (b) NISTCHO cell lines stained with plant-derived lectins; (c) CHO VRC01 cell lines stained with RPLs; and (d) NISTCHO cell lines stained with RPLs. **Note:** The NISTCHO parental sample stained with SNA was excluded from quantitative analysis in panel (b) due to the absence of detectable cells and fluorescence signal under plate-based confocal imaging conditions. Although slide-based imaging revealed low-level background fluorescence and surface lectin binding, differences in sample preparation and imaging modality precluded direct comparison with plate-based datasets. Consequently, this condition was omitted to ensure consistency and comparability across samples.

### Decoupling of Surface and Product Glycosylation for β4GALT1 and ST3GAL4

In contrast to FUT8 and COSMC, perturbation of β4GALT1 and ST3GAL4 did not produce clearly distinguishable surface glycosylation phenotypes despite their pronounced effects on antibody glycosylation. Neither lectin-aided flow cytometry nor confocal imaging reliably differentiated β4GALT1 knockout or overexpression clones from parental controls using galactose-binding lectins. Similarly, ST3GAL4 knockout cells retained detectable α2,3-sialylation at the membrane surface despite producing antibodies devoid of α2,3-sialylated glycans (Figures 2c–f).

These findings are consistent with extensive functional redundancy within the β4GALT and ST3GAL enzyme families in CHO cells, which can sustain surface glycan presentation even when a dominant isoenzyme is disrupted [15–17]. Importantly, while confocal imaging revealed spatially localised lectin binding following these perturbations, flow cytometry demonstrated a net reduction in global surface glycan abundance. As a result, lectin-assisted enrichment was ineffective for these targets, necessitating clone isolation by limiting dilution followed by molecular validation. Importantly, this decoupling underscores that surface glycosylation cannot be assumed to mirror product glycosylation for all glycogenes.

### Comparison of Plant-Derived and Recombinant Lectins for Surface Glycan Detection

To evaluate the robustness of lectin-based profiling, plant-derived lectins were compared with recombinant prokaryotic lectins (RPLs) across engineered cell lines. While both lectin classes detected expected trends in the CHO VRC01 background, RPL staining exhibited greater variability and broader fluorescence distributions, particularly in NISTCHO cells. This increased variability likely reflects additional processing steps required for RPL labelling and reduced binding consistency under fixed-cell imaging conditions.

Overall, pre-conjugated plant lectins provided more reproducible and interpretable measurements of global surface glycan abundance by flow cytometry, whereas confocal imaging offered complementary spatial information regarding glycan distribution rather than absolute surface load. In addition, recombinant lectins may be better suited to live-cell applications where cytotoxicity is a concern (Figure 3).

## Discussion

This study highlights a central challenge in CHO glycoengineering: genetic modifications that strongly alter product glycosylation do not necessarily produce resolvable changes at the cell surface. The degree of concordance between surface and product glycosylation is highly dependent on the targeted enzyme, pathway architecture, and host-cell background [13].

Edits targeting enzymes with limited redundancy, such as COSMC and FUT8, produced clear and consistent surface phenotypes that closely mirrored product glycosylation [14, 18]. In contrast, perturbations involving highly redundant pathways, notably β4GALT1 and ST3GAL4, resulted in pronounced changes in antibody glycosylation while leaving the surface glycome largely intact [15, 17, 19]. This redundancy arises from the presence of multiple β4GALT and ST3GAL isoenzymes in CHO cells. Importantly, emerging evidence suggests that β4GALT1 exhibits preferential activity toward Fc glycan processing, such that its disruption disproportionately affects mAb galactosylation, while other β4GALT isoenzymes continue to support glycosylation of endogenous cell-surface glycoproteins [15]. This division of labour between isoenzymes likely underpins the reduced correlation observed between product and surface galactosylation. Collectively, these findings underscore the buffering capacity of the CHO surface glycosylation network and explain why lectin-based sorting and screening fail for certain targets embedded within redundant, substrate-diverse pathways despite successful modulation of product glycosylation.

Although both lectin-assisted flow cytometry and confocal microscopy rely on labelling of cell-surface glycans, the two approaches report distinct aspects of glycan presentation. Flow cytometry integrates fluorescence across the entire plasma membrane and provides a population-averaged signal of total surface lectin binding, while confocal microscopy imaging detects spatial organisation and accessible glycan microdomains. This distinction is important as lectin binding is context-dependent – lectins can recognise overlapping and heterogeneous glycan motifs on different glycoproteins, and binding patterns heavily depend on accessibility and local distribution on the cell surface rather than merely abundance alone [20, 21]. This is particularly relevant for perturbations of β4GALT1 and ST3GAL4, where the presence of multiple isoenzymes with differing substrate specificities enables continued glycosylation of distinct protein pools. As a result, disruption of a dominant isoenzyme can markedly alter mAb glycosylation while preserving glycan presentation on subsets of cell-surface glycoproteins, leading to an apparent decoupling between global abundance and spatially resolved lectin binding. Consequently, residual lectin staining detected by confocal imaging does not necessarily indicate preservation of product-level glycosylation.

Importantly, gain-of-function engineering via ST6GAL1 overexpression emerged as a distinct class of intervention. By introducing a non-native enzymatic activity, this modification generated concordant and readily detectable surface and product phenotypes [17, 19, 22]. This suggests that lectin-based assays are particularly well suited for validating single pathway additions, while being less reliable for loss-of-function edits within endogenous, redundant networks.

Differences observed between VRC01 and NISTCHO further demonstrate that even closely related CHO lineages cannot be assumed to share identical surface glycomes. Baseline host-cell differences influence both lectin binding and the phenotypic manifestation of engineered changes, reinforcing the need for host-specific characterisation.

From a methodological perspective, lectin-assisted confocal microscopy and flow cytometry should be viewed as complementary tools. Flow cytometry offers rapid, high-throughput assessment when strong surface phenotypes are present, whereas confocal microscopy provides critical spatial and qualitative insight when phenotypes are subtle or ambiguous [23, 24]. Plant-derived lectins, supplied pre-conjugated, were more reliable for fixed-cell imaging in this study. However, such lectins may introduce variability due to impurities or cytotoxic effects that can compromise cell integrity. In contrast, recombinant lectins, while generally exhibiting lower binding specificity, are less prone to inducing cellular damage and may therefore be better suited for live-cell applications and lectin-based enrichment strategies, provided their specificity aligns with the target glycan[25, 26].

In conclusion, lectin-based surface profiling is a valuable component of CHO glycoengineering workflows, but its interpretability is inherently context-dependent. Surface glycosylation should not be used as a universal surrogate for product glycosylation. Instead, lectin assays are most effective when applied with prior knowledge of enzyme redundancy, host-cell background, and the expected nature of the engineered modification. In this context, lectin-based readouts can be particularly valuable during early pool screening or micro-pool enrichment, providing a rapid, low-complexity means of identifying pools enriched for the desired glycoengineering outcome [27]. This makes them attractive alternatives to extended single-cell cloning workflows, helping to reduce development timelines and aligning with upstream screening strategies increasingly explored in industrial settings, particularly when prioritising speed and throughput over absolute specificity [28, 29]. When used judiciously and in combination with direct product analysis, these tools can accelerate clone characterisation and support rational design of glycoengineered CHO production platforms.

## Methods

### Plasmid Construction

CRISPR/Cas9-mediated gene knockouts were performed using pSpCas9(BB)-2A-GFP (pX458, Addgene #48138) [30]. The sgRNAs targeting Fut8, β4GalT1, COSMC, and ST3Gal4 were cloned under the U6 promoter using BpiI sites [14, 31]. The sgRNA sequences and genomic target sites are provided in Supplementary Table S1 and Figure S1. The ST6Gal1 overexpression plasmid was purchased from Invivogen (puno1-hst6gal1a) and selection conducted with blasticidin (Invivogen, Cat. No. ant-bl-1).

### Cell Culture and Transfection

CHO VRC01 and NISTCHO cell lines were cultured under standard suspension conditions in chemically defined media: (i) ActiPro™ cell culture media (HyClone™, Cat. No. SH31039.02) supplemented with 4 mM L-glutamine (Gibco™, Cat. No. 25030-024) and 100 nM methotrexate (MTX); and (ii) EX-CELL® CD-CHO Fusion Medium (SAFC, Cat. No. 14365C/24365C) without L-glutamine, respectively. Cells were maintained at 37°C, 5% CO2, and humidified air, and passaged every 2–3 days. Plasmids were delivered by electroporation using the Amaxa® Cell Line Nucleofector® Kit V (Lonza Bioscience, Cat. No. VCA-1003). More details on the transfection procedure can be found in the Supplementary Information. Post-transfection recovery was performed in pre-warmed complete media prior to downstream selection.

### Clone Isolation and Validation

Edited populations were enriched by lectin-aided fluorescence-activated cell sorting (FACS). FUT8 and COSMC knockouts, and ST6GAL1 overexpression were identified by lectin staining, while β4GALT1 and ST3GAL4 knockouts were isolated by limiting dilution and confirmed by deletion PCR using primers listed in Supplementary Table S2.

### Antibody Purification and Glycan Analysis

Monoclonal antibodies were purified from day-7 culture supernatants using an ÄKTA™ Start purification system (Cytiva Life Sciences) equipped with a HiTrap™ Protein A HP column (GE Healthcare), according to the manufacturer’s instructions.

For all samples, intact native mass spectrometry was employed to enable comprehensive assessment of sialylated glycoforms across all N-glycosylation sites. Briefly, 10 µg of each purified antibody sample was injected onto a Vanquish™ Flex UHPLC system hyphenated to a Thermo Scientific™ Exploris™ MX mass spectrometer operated in intact protein mode at a resolving power of 30,000 (at m/z 200). Online buffer exchange was performed using a NativePac™ OBE-1 size-exclusion chromatography column (2.1 x 50 mm, 3 µm particle size, Thermo Scientific) under isocratic conditions with 50mM ammonium acetate at a flow rate of 0.2 mL/min.

Raw spectra corresponding to the intact antibody elution peak were deconvoluted using BioPharma Finder™ software (version 5.3, Thermo Scientific) employing the ReSpect algorithm with default native analysis settings. Deconvoluted masses were matched against the VRC01 or NIST mAb protein sequence, allowing up to four or two variable N-glycosylation modifications corresponding to the expected glycosylation sites. Pyroglutamate formation at the heavy-chain N-terminus and C-terminal lysine clipping were specified as fixed modifications for both proteins. Fucosylation, galactosylation, and sialylation indices were calculated from relative glycoform abundances, using Equations 1 to 3. Where *v*_*fuc or gal or sia*,*i*_ is the number of fucose, galactose or sialic acid residues in glycan *i, x*_*i*_ is the relative abundance of glycan *ii*, and *v*_*gal or sia*,*max*_ is the maximum number of galactose or sialic acid residues that the glycan can contain (e.g., *v* = 2 for biantennary glycans).

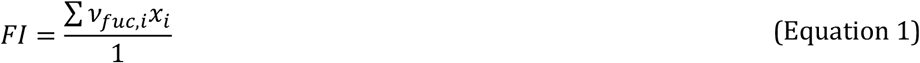

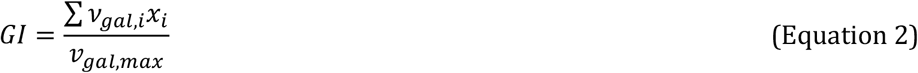

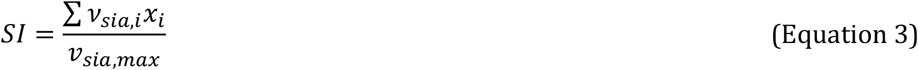

### Lectin-Aided Flow Cytometry

Cell-surface glycosylation was quantified by flow cytometry on day 3 of cell culture using plant-derived and recombinant prokaryotic lectins. Autofluorescence-corrected median fluorescence intensity (MFI) and staining index (SI) were calculated to normalise background differences between cell lines (Equations 4 and 5). Detailed procedures for sample preparation, MFI and SI calculations are described in the Supplementary Information. Flow cytometry data (FCS files) were gated using the CytExpert v.2.4 software (Beckman Coulter). Statistical analysis was performed using Excel with significance assessed by two-tailed *t*-tests and a threshold of *p* < 0.05. Statistical significance was denoted as: **** *p* < 0.0001, *** *p* < 0.001, ** *p* < 0.01, * *p* < 0.05, and ns (not significant) for *p* ≥ 0.05.

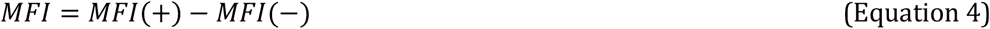

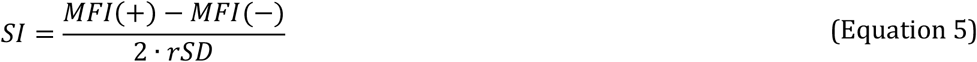

### Confocal Microscopy Sample Preparation and Staining

All confocal microscopy images were acquired using a LSM900 (Zeiss, Germany) equipped with a Plan-Apochromat 20x/0.8 NA objective to visualise and qualitatively validate lectin binding patterns on engineered CHO cell lines. Cells were seeded at 5,000–20,000 cells per well in 96-well ibiTreat µ-plates with #1.5 polymer coverslip bottoms (Ibidi GmbH, Cat. No. 89606) and incubated overnight. Cells were fixed with 4% paraformaldehyde, blocked with bovine serum albumin (BSA), and stained with recombinant lectins. Following washing, samples were imaged directly in the polymer-bottom plates.

### Image Analysis and Quantification

Automated image analysis was performed using ImageJ/Fiji (v1.54). Confocal Z-stacks (16-bit) were imported via the Bio-Formats plugin to maintain spatial scaling. For each stack, a 32-bit Sum Intensity Projection was generated to capture the total fluorescence signal across the entire cellular volume. To correct for non-specific background and uneven illumination, a rolling-ball background subtraction (radius = 50 μm) was applied to the projections.

Cell segmentation was achieved by creating a binary mask from the processed projections. The images were pre-filtered with a median filter (radius = 2) and a Gaussian blur (σ = 1) to reduce pixel noise. Automated thresholding was performed using the Triangle algorithm, followed by “Fill Holes” and “Dilate” operations to ensure a solid mask covering the full extent of the membrane-stained cells. Touching objects were separated using a Watershed transform.

Quantitative measurements were obtained using the “Analyse Particles” tool, targeting objects with an area of 100 – 700 μm^2^ and a circularity of 0.35 – 1.00. Objects touching the image edge were excluded to ensure measurement accuracy. To account for variations in Z-stack depth across samples, the mean fluorescence intensity (MFI) of each identified cell was normalised by dividing the total intensity by the number of focal planes (Z-slices) within that specific stack. This “Mean per Slice” value was used for all subsequent statistical comparisons and plotting.

### Statistical Analysis for Confocal Microscopy Assay

Statistical analysis was performed in R (version 4.5.2). Normality of fluorescence intensity distributions was assessed using the Shapiro-Wilk test for each sample group. As the majority of groups exhibited non-normal distributions (*p* < 0.05), non-parametric statistical testing was employed. Pairwise comparisons between parental and modified cell lines were performed using the Wilcoxon rank-sum test (Mann-Whitney U test) for each plant-derived or recombinant prokaryotic lectin independently. Statistical significance was denoted as: **** *p* < 0.0001, *** *p* < 0.001, ** *p* < 0.01, * *p* < 0.05, and ns (not significant) for *p* ≥ 0.05.

## Acknowledgments

The authors gratefully acknowledge funding from: (i) Research Ireland (Grant Numbers 20/FFP-P/8864 and 18/EPSRC-CDT/3587); (ii) the UKRI Engineering and Physical Sciences Research Council (Grant No. 18/EPSRC-CDT/3587); and (iii) the European Research Council (ERC) (Grant No. 101052376). The authors also thank Robert Dunne from GlycoSelect for supplying the RPLs used in this study and UCD Conway Flow Cytometry Core Facility for access to the flow cytometers for the analyses. The graphical abstract was created with images from Flaticon (https://www.flaticon.com/).

## Author Contributions

C.A.R., S.L.Y.L., and I.J.V conceived and designed the study. C.A.R. and S.L.Y.L. generated the genetic constructs and cell lines, produced mAb samples, performed flow cytometry analysis, and prepared samples for confocal microscopy. J.B. performed confocal microscopy analysis. S.C. performed product glycan analysis. S.L.Y.L and C.A.R. wrote the manuscript. J.B., E.C., and I.J.V provided supervision and acquired funding. All authors reviewed and approved the manuscript.

