## Supplementary Information for "Unmasking Glycoforms: Lectin-Based Profiling and Functional Implications of Targeted Glycosylation Knockouts in CHO Cells"

Cristina Abascal Ruiz[1], Sheryl Li Yan Lim[1], Jacobus Brink[1], Sara Carillo[2], Eoin Casey[1], Jonathan Bones[1, 2], and Ioscani Jiménez del Val[1,*]

[1] School of Chemical & Bioprocess Engineering, University College Dublin, D04 V1W8, Ireland;

[2] National Institute for Bioprocessing Research & Training, Characterisation and Comparability Laboratory, A94 X099, Ireland

*Author to whom correspondence should be addressed.

### Methods

#### Electroporation

Two recombinant CHO cell lines were used: (i) CHO VRC01 and (ii) NISTCHO. CHO VRC01 is derived from CHO-K1 and expresses a broadly neutralising IgG1κ antibody targeting HIV-1 CD4 binding sites, which contains four glycosylation sites – two in the Fc and two in the Fab fragment. NISTCHO is a clonal CHO-K1 line producing the cNISTmAb reference antibody and contains two consensus IgG1 glycosylation sites in the Fc fragment. Viable cell density and viability assessed using a Countess™ 3 FL Automated Cell Counter.

Electroporation was performed using the Amaxa® Cell Line Nucleofector® Kit V. Cells ($2.5 \times 10^6$ cells per cuvette) were pelleted (400×g for CHO VRC01 or 200×g for NISTCHO, 8 min) and resuspended in 82 µL Nucleofector solution and 18 µL supplement containing up to 5 µg plasmid DNA. Electroporation was carried out using programme “Cell 9,” followed by a second identical pulse after a 2-minute rest. Cells were allowed to recover briefly before transfer into pre-warmed medium and incubated for 24 h prior to downstream selection by FACS or limiting dilution, as appropriate.

### Lectin Preparation

Two sources of lectins were used in this study, and each preparation workflow are detailed below.

#### *Plant-Derived Lectins*

Plant lectins used included Aleuria aurantia lectin (AAL, FL-1391-2), Vicia villosa lectin (VVL, FL-1231-2), Maackia amurensis lectin I (MAL I, FL-1311-2), Erythrina cristagalli lectin (ECL, FL-1141-5), and Sambucus nigra agglutinin (SNA, B-1305), each recognising distinct glycan motifs relevant to antibody glycosylation. All lectins were sourced from Vector Laboratories; AAL, VVL, MAL I, and ECL were supplied FITC-conjugated, while SNA was Cy3-conjugated.

Cells were incubated directly with lectins at a final concentration of 10 μg/mL for 15 minutes at room temperature in darkness. Samples were washed three times with PBS prior to flow cytometric or imaging analysis.

#### *Recombinant Prokaryotic Lectins (RPLs)*

Recombinant prokaryotic lectins (RPL-Fuc1, RPL-Gal4, and RPL-Sia1) were used to detect α-linked fucose, terminal β-galactose/LacNAc, and α2-3-linked sialic acid, respectively. The lectins were sourced from GlycoSeLect (Ireland; L-011-2 mg, L-005-2 mg, L-008-2 mg, respectively). As these lectins were supplied unlabelled, biotinylation was performed using NHS-PEG4-Biotin (Thermo Scientific, Cat. No. A39259) followed by fluorescent conjugation with DyLight™ 488-streptavidin (Invitrogen™, Cat. No. 21832).

Briefly, 10 μM RPL was incubated with 40 μM NHS-PEG4-Biotin on ice for 2 h, and excess biotin was removed by desalting. Biotinylated lectins (8 μg/mL) were conjugated to streptavidin-DyLight™ 488 and incubated with $0.5 \times 10^6$ washed cells for 30 minutes at room temperature in TBS containing 1 mM of $Ca^{2+}$, $Mg^{2+}$, and $Mn^{2+}$. After washing, samples were resuspended in TBS and analysed.

### Figures and Tables

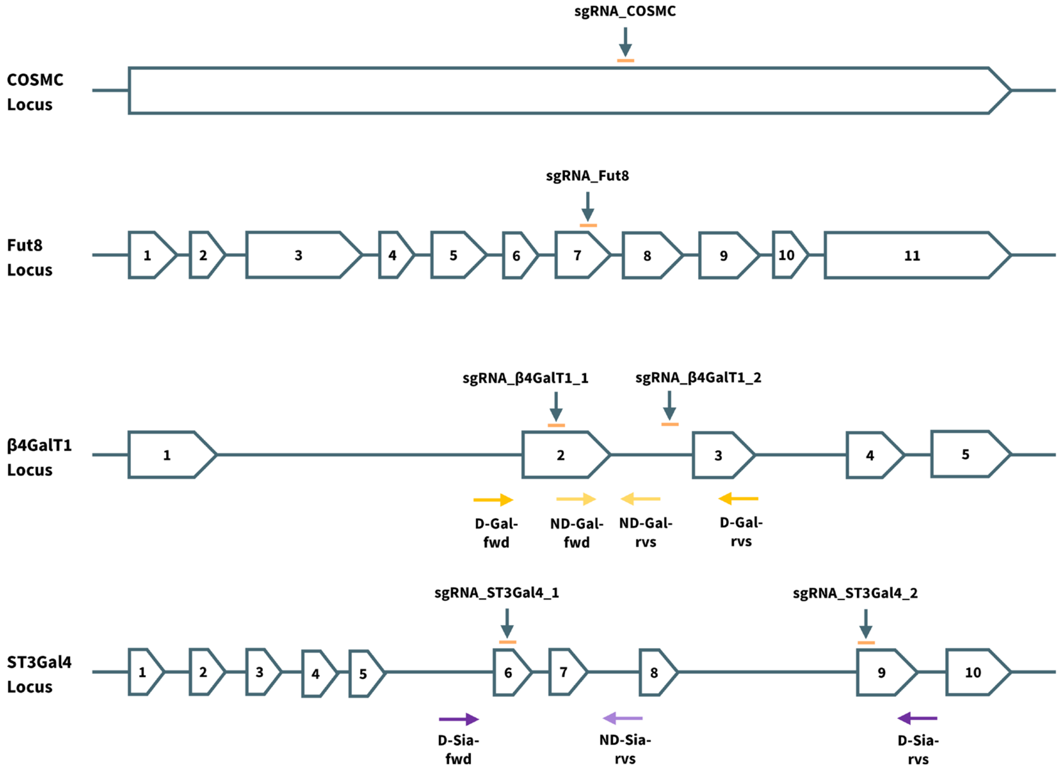


Supplementary Figure S1: CRISPR/Cas9 mediated knockouts of endogenous COSMC, FUT8, β4GALT1, and ST3GAL4 genes, and the associated non-deletion/deletion PCR primer binding sites on the loci

Supplementary Table S1: sgRNA sequences for CRISPR/Cas9 knockout

| Knockout | Name | Sequence (5'→3') | PAM (5'→3') |
|---|---|---|---|
| **COSMC** | sgRNA_COSMC | GAATATGTGAGTGTGGATGG | AGG |
| **FUT8** | sgRNA_FUT8 | GATCCGTCCACAACCTTGGC | TGG |
| **β4GALT1** | sgRNA_βGALT1_1 | ATTGCAACAGAAATGTGCCG | GGG |
| | sgRNA_βGALT1_2 | TAGTGAGTCAGACCAAGACG | GGG |
| **ST3GAL4** | sgRNA_ST3GAL4_1 | CGGTTGCGGAACAGTTCACT | GGG |
| | sgRNA_ST3GAL4_2 | GATCACGCTTAAGTCTATGG | CGG |

Supplementary Table S2: Non-deletion/deletion primer sequences

| Knockout | Name | Sequence (5'–3') (Binding Regions Bolded) |
|---|---|---|
| **β4GALT1** | ND-Gal-fwd | AATGTAGTTCTTCTTGAGTGCC |
| | ND-Gal-rvs | AAATACAGAACTGGGATCTGAA |
| | D-Gal-fwd | CTGGAAATGGATTGTTGACTCAGAGGG |
| | D-Gal-rvs | GAGAACCATCACATAAACTAAGGAAAACACC |
| **ST3GAL4** | ND-Sia-rvs | TCTCCTCCTAGCTTGGTCTCAC |
| | D-Sia-fwd | GGGCTGAGGAGAACCTAGAAGT |
| | D-Sia-rvs | CCACTTCTGACACTGAATGACC |
